# DNA methylation reprogramming in marsupial embryos is restricted to the extraembryonic lineage

**DOI:** 10.1101/2025.11.09.686659

**Authors:** Allegra Angeloni, Jillian M. Hammond, Timothy J. Peters, Andre L. M. Reis, Leah Kemp, Timothy Amos, Hasindu Gamaarachchi, Sam Humphries, Lynda A. Wilmott, Suranjana Pal, V. Pragathi Masamsetti, Megan Weatherstone, Kenny Chi Kin Ip, Karina Pazaky, Alice Steel, Ruth Lyons, Elly D. Walters, Ning Liu, Patrick Tam, Jose M. Polo, Paul Waters, Susan J. Clark, Linda J. Richards, Andrew D. Smith, Heather Lee, Ira W. Deveson, Oliver W. Griffith, Ksenia Skvortsova

## Abstract

DNA methylation (5mC) is an epigenetic mark that plays a critical role in defining cell fate. Following fertilisation, DNA methylation inherited from gametes must be reprogrammed to establish totipotency and enable the parental-to-zygotic transition. To accomplish this, non-mammalian vertebrates such as zebrafish and medaka subtly reprogram maternal 5mC profiles while maintaining high methylation levels throughout embryogenesis. In contrast, eutherian mammals such as mouse and human undergo global 5mC erasure in both embryonic and extraembryonic lineages. However, while embryonic 5mC is rapidly re-established to high levels upon implantation, the trophectoderm, which gives rise to the placenta, displays sustained and conserved DNA hypomethylation, suggesting that this drastic 5mC erasure may be functionally linked to complex placentation in mammals. To clarify whether extensive post-fertilisation 5mC erasure co-evolved with placentation, we explored embryonic methylation dynamics in marsupials, a lineage of therian mammals with a short-lived placenta. We produced a near complete telomere-to-telomere (T2T) genome and generated detailed epigenome maps of embryonic development for an Australian marsupial, the fat-tailed dunnart (*Sminthopsis crassicaudata*). We found the dunnart embryo exhibits genome wide DNA demethylation at the blastocyst stage, but these changes occur in the trophectoderm only, suggesting that 5mC erasure in the placenta is an ancestral state in therian mammals. Furthermore, the T2T-level dunnart genome assembly enabled identification of sex chromosomes, uncovering extensive hypomethylation of the paternally-inherited inactive X chromosome in females and revealing the previously unannotated master regulator of X chromosome inactivation, lncRNA *Rsx*. Our data indicate that while the use of genome-wide 5mC erasure differs between eutherian and marsupial lineages, 5mC erasure in extraembryonic tissue is ancestral to therian mammals and may be necessary to support placental development.

**HIGHLIGHTS:**

- First embryonic DNA methylation maps in an Australian marsupial
- Extensive global erasure of DNA methylation in the trophectoderm
- Maintenance of high DNA methylation in the embryonic lineage
- Hypomethylated paternal X chromosome with methylated escapee genes

## MAIN

DNA methylation at cytosine residues (5-methylcytosine, 5mC) is the most prevalent DNA base modification in eukaryotic genomes, possessing crucial roles in gene regulation^1,2^. During early vertebrate development, 5mC landscapes inherited from both parents must be co-ordinately reprogrammed to establish epigenome equivalence^3^. This 5mC reprogramming in the early embryo is hypothesised to be associated with the establishment of totipotency and proper embryonic genome activation, with compromised 5mC reprogramming being tightly linked to developmental arrests in cloned embryos^4^.

In eutherian mammals, such as mice and humans, 5mC is globally erased during two critical developmental windows: first immediately following fertilisation^5–9^, and next during primordial germ cell development^10–14^, with an additional demethylation event occurring during spermatogenesis in males^15,16^. Post-fertilisation reprogramming involves active DNA demethylation of the paternal genome in the pronucleus, followed by passive DNA demethylation of both maternal and paternal genomes during cleavage stages^6,7,9,17–19^. Intriguingly, studies in non-mammalian vertebrates have revealed strikingly different strategies for managing inheritance and reprogramming of parental epigenomes^20–25^. For example, in zebrafish, medaka, and sea lamprey, the paternal 5mC pattern is largely maintained throughout early embryogenesis, while the maternal genome undergoes reprogramming to match the paternal methylome prior to zygotic genome activation^20–22,24,25^. This results in inheritance rather than erasure of the sperm methylation pattern, which is fundamentally different from the mammalian paradigm.

These phylogenetic differences in 5mC reprogramming strategies within the vertebrate phylum raise fundamental questions about the evolutionary origins and functional significance of the extensive demethylation observed in most eutherian mammals^26^. Given that non-mammalian vertebrates accomplish development without drastic 5mC erasure, the extensive reprogramming in eutherians may have evolved to support mammalian-specific developmental innovations, such as early specification of extraembryonic lineages and complex uterine implantation processes^27–30^. The formation of trophectoderm (TE) and inner cell mass (ICM) represents the first cell fate specification event during eutherian embryogenesis. TE cells become lineage-restricted to trophoblast derivatives that are essential for placentation, whereas ICM cells lose trophoblast potential but retain pluripotency to generate all cell types of the embryo proper as well as primitive endoderm and extraembryonic mesoderm^31,32^. Importantly, whereas the embryonic genome is rapidly remethylated to high levels upon implantation, trophectoderm derivatives maintain prolonged DNA hypomethylation, suggesting a key role for hypomethylation in placenta development^33–35^. The unique post-fertilisation erasure of 5mC in both embryonic and extraembryonic lineages in eutherian mammals, which is entirely absent in non-mammalian vertebrates studied to date, supports a causal relationship between 5mC reprogramming and the emergence of placentation. However, determining whether extensive 5mC erasure is unique to eutherian mammals or also occurs in other mammalian lineages is critical for understanding its evolutionary significance.

Marsupials represent an independent lineage of live-bearing mammals that diverged from eutherians ∼160 million years ago. Marsupials possess unique developmental traits, including altricial young and a short gestation with a brief period of placentation^36,37^. Despite their short gestation, marsupials have a functional placenta that provides all embryonic nourishment prior to birth, formed through the interaction of the trophectoderm and the uterine mucosa^36^. In many marsupials, the trophectoderm invades through the uterine epithelium to sit adjacent to endometrial blood vessels. Given their unique evolutionary position, marsupials allow us to understand whether 5mC erasure is a unique feature of eutherian mammals, or if it evolved prior to the evolution of a placenta in therian mammals. While methylation associated with genomic imprinting has been studied widely in marsupials^38–46^, our current understanding of embryonic 5mC reprogramming in non-eutherian mammals is extremely limited, with only two studies focused on the South American opossum (*Monodelphis domestica*), being published to date^47,48^. The Australian fat-tailed dunnart (*Sminthopsis crassicaudata*) has recently emerged as a model marsupial species for comparative developmental biology, owing to their relative ease of husbandry, well-characterised embryogenesis, post-natal development, and unique placental biology^49^. Critically however, the gene regulatory mechanisms governing embryonic development in the dunnart remain largely uncharacterised, unlike in the American opossum model. Given the ∼80-million year phylogenetic divergence between Australian and American marsupials, investigating these mechanisms in the dunnart is pertinent to understanding the divergence and conservation of developmental programs across the mammalian lineage.

Here, we present a near-complete telomere-to-telomere (T2T) genome assembly and the first comprehensive genome-wide 5mC profiling of embryonic development in an Australian marsupial, the fat-tailed dunnart (*Sminthopsis crassicaudata*). While our findings reveal retention of a hypermethylated genome in the fat-tailed dunnart embryo that is distinct from what is described in eutherians, the dunnart undergoes eutherian-like extensive 5mC erasure in the trophectoderm. These dynamics mirror developmental 5mC patterns described in the opossum^47,48^, suggesting a deeply conserved role for DNA hypomethylation in placental development and regulation across evolutionarily diverse mammalian lineages. Additionally, generation and analysis of T2T, gap-free sex chromosomes identified the previously unannotated long non-coding RNA *Rsx,* which is responsible for silencing the paternal X chromosome in marsupials^50^; as well as global hypomethylation of the inactive X in the female dunnart, with the exception of genes that escape inactivation. Our findings present the first detailed exploration of embryonic 5mC dynamics in the fat-tailed dunnart, providing new insights into the functionality of developmental epigenome reprogramming strategies across diverse mammalian clades.

## RESULTS

### Genome-wide DNA methylation reprogramming in marsupial embryos

First, to improve upon the existing scaffold-level fat-tailed dunnart genome^51^, we generated a near-complete telomere-to-telomere (T2T) assembly. To achieve this, we combined long-read Oxford Nanopore Technology (ONT)^52^ with chromosome conformation capture (Hi-C) data^53^, and employed the novel ‘Cornetto’ adaptive assembly strategy^54^. The resulting genome assembly identified six autosomes and a pair of sex chromosomes (X and Y), representing a significant improvement in assembly contiguity with an N50 of 669Mb, nearly an order of magnitude greater than the 79Mb N50 of the published genome (Fig. 1A-D). We achieved complete gap-free, telomere-to-telomere (T2T) contig assemblies for chromosomes X, Y, and 6. Chromosome 1 was assembled as a T2T scaffold, and chromosomes 2-5 were near-complete (Fig. 1D). Notably, the 12.1 Mb Y chromosome identified in our assembly corroborates earlier cytogenetic evidence that the fat-tailed dunnart harbors the smallest documented mammalian Y chromosome, representing only 20% of human Y chromosome length^55^. This T2T assembly provided a robust reference for DNA methylation analyses.

**Figure 1:**
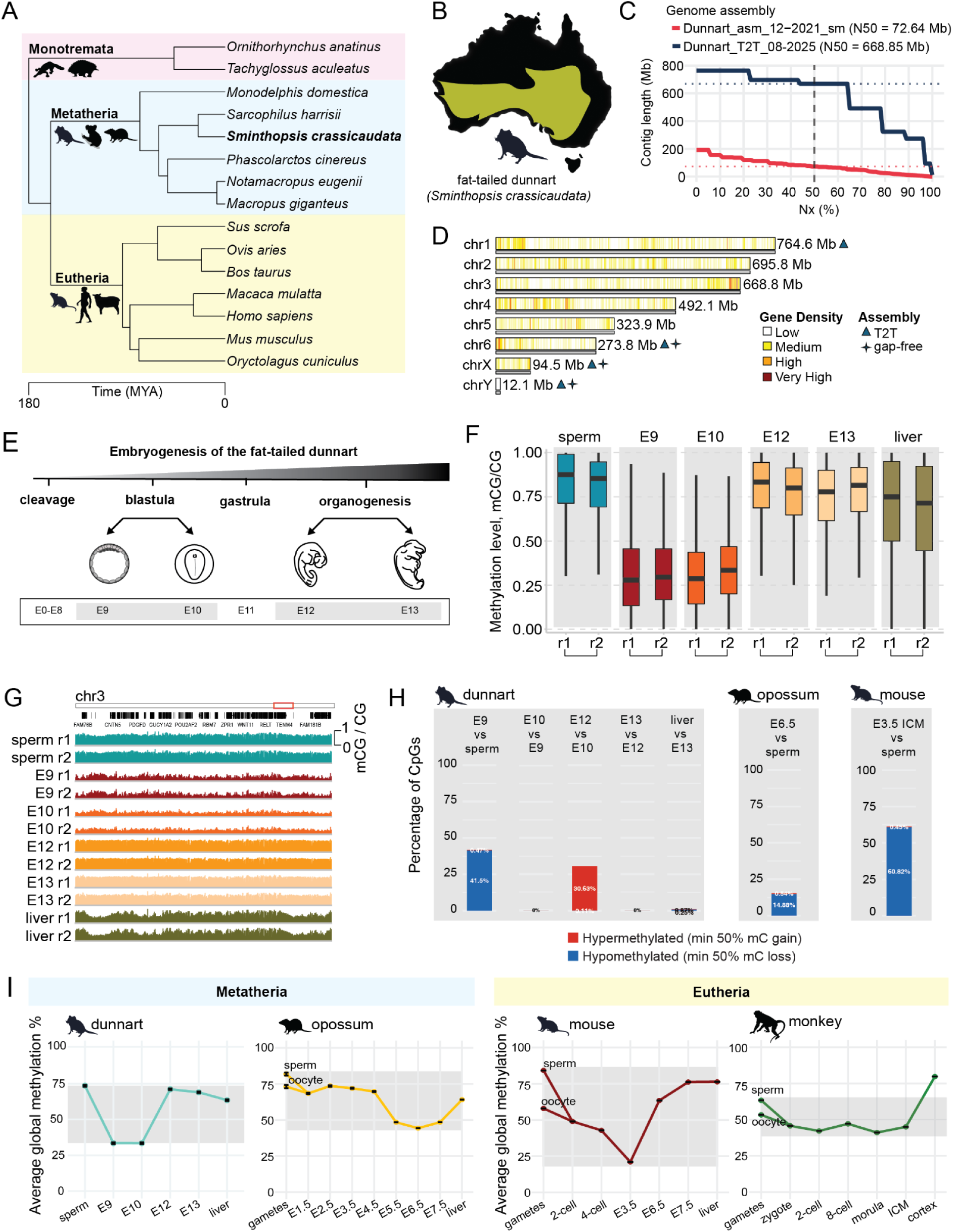
Global DNA methylation dynamics in the fat-tailed dunnart embryo. **(A)** Phylogenetic relationships of the monotremes, meta- and eutherian mammals^59^. **(B)** Map of Australia shows the habitat of the fat-tailed dunnart (*Sminthopsis crassicaudata*). **(C)** Nx plot displaying contigs ordered by length against cumulative genome coverage, comparing the current publicly available assembly (red) with the assembly generated in this study (blue)^51^. **(D)** Chromosome level assembly of the fat-tailed dunnart genome displaying chromosome sizes and gene density. **(E)** Schematic representation of the development stages of the fat-tailed dunnart embryo. Embryo drawings adapted^49^. **(F)** Boxplots of genomic 5mC levels (1 kb non-overlapping genomic tiles) of sperm, embryonic (E9, E10, E12, E13) and adult somatic tissue (liver) in two biological replicates (r1 and r2). The boxes show the interquartile range (IQR) around the median. The upper and lower whiskers extend from the hinge to the largest and smallest value, respectively, no further than 1.5 IQR. **(G)** IGV browser track depicting DNA methylome profiles of fat-tailed dunnart sperm, four embryonic stages, and adult liver tissue. **(H)** Percentages of differentially methylated CpGs in the genome between consecutive stages of embryonic development in the dunnart, opossum and mouse. CpGs with at least 50% methylation loss compared to the previous stage were considered as hypomethylated. CpGs with at least 50% methylation gain compared to the previous stage were considered as hypermethylated. **(I)** Line plots depicting global DNA methylation dynamics (mean mC value across all CpGs in the genome) in the early embryo of dunnart (*Sminthopsis crassicaudata*), opossum (*Monodelphis domestica*), mouse (*Mus musculus*) and monkey (*Macaca mulatta*)^17,48,57,60^.

To examine post-fertilisation DNA methylation dynamics in the fat-tailed dunnart, we collected sperm, embryos and adult tissue. Embryonic samples spanned four developmental stages: 9 days post-fertilisation (E9, stage 17 - bilaminar blastocyst), 10 days post-fertilisation (E10, stage 19 - trilaminar blastocyst), 12 days post-fertilisation (E12, stage 28 - late organogenesis), and 13 days post-fertilisation (E13, stage 31 - implantation), with biological replicates for each stage (Fig. 1E)^49^. Single-base resolution DNA methylation profiling was performed using enzymatic methyl-seq^56^ (Supplementary Table 1). Principal component analysis (PCA) identified separation of E9 and E10 methylomes from other samples (Supplementary Fig. 1A). Analysis of DNA methylation per 1kb genomic tiles revealed substantial erasure of gametic DNA methylation in the blastocyst at E9 and E10 (Fig. 1F, G) with only a small fraction of CpG sites retaining high (0.8-1.0) methylation levels (Supplementary Fig. 1B). Differential DNA methylation analysis revealed ∼42% of all CpG sites losing at least 50% methylation in the E9 blastocyst compared to sperm (Fig. 1H) and ∼53% of all CpG sites losing at least 20% methylation (Supplementary Fig. 1C). This erasure of DNA methylation was more extensive in the dunnart blastocyst than the opossum blastocyst, but less than the mouse blastocyst (Fig. 1H, Supplementary Fig. 1C). Finally, to compare DNA methylation dynamics between metatherian and eutherian mammals, we plotted average methylation levels during early embryonic development encompassing the blastocyst stage of dunnart, opossum, mouse and rhesus monkey^17,48,57^. The analysis revealed that while sperm methylation levels are consistently high (>70%) across all these species, embryonic methylomes reach different minimum levels depending on the species. Mouse embryos show the most extensive demethylation, dropping to 21% (E3.5), followed by dunnart at 33% (E9/E10), monkey at 41% (morula), and opossum at 44% (E6.5) (Fig. 1I, Supplementary Table 2).

Collectively, this demonstrates the substantial DNA methylation erasure in the dunnart’s bi- and trilaminar blastocyst, but highlights the varying levels of DNA methylation reprogramming across mammals, with notable differences even between eutherians^58^.

### DNA methylation is largely retained in the embryonic lineage but erased in the trophectoderm

To explore genomic regions that undergo 5mC erasure, we examined differentially methylated regions between consecutive developmental stages^61^. Specifically, we focussed on DMRs between sperm and E9 to determine early hypomethylated regions; and between E9 and E12 to determine late hypomethylated regions. Based on the average genomic 5mC levels, the majority of hypomethylated DMRs were those present between sperm and E9 (early hypoDMRs, n=199,254), while only a small subset of regions showed hypomethylation at E12 (late hypoDMRs, n=83) (Fig. 2A). While late hypoDMRs showed significant enrichment over promoters and exons, early hypoDMRs were more uniformly distributed across the genome suggestive of a genome-wide DNA demethylation in the E9 embryo (Fig. 2B). However, given that EM-seq was performed on whole embryos, it remained to be determined whether the extent of DNA demethylation differs between embryonic and extraembryonic lineages of the blastocyst. To determine in which lineage DNA methylation reprogramming takes place, we sorted single nuclei from E9 dunnart embryos and performed DNA methylation profiling using single cell bisulfite sequencing (scBS-seq)^62^, generating 86 high quality methylomes (Supplementary Table 3).

**Figure 2:**
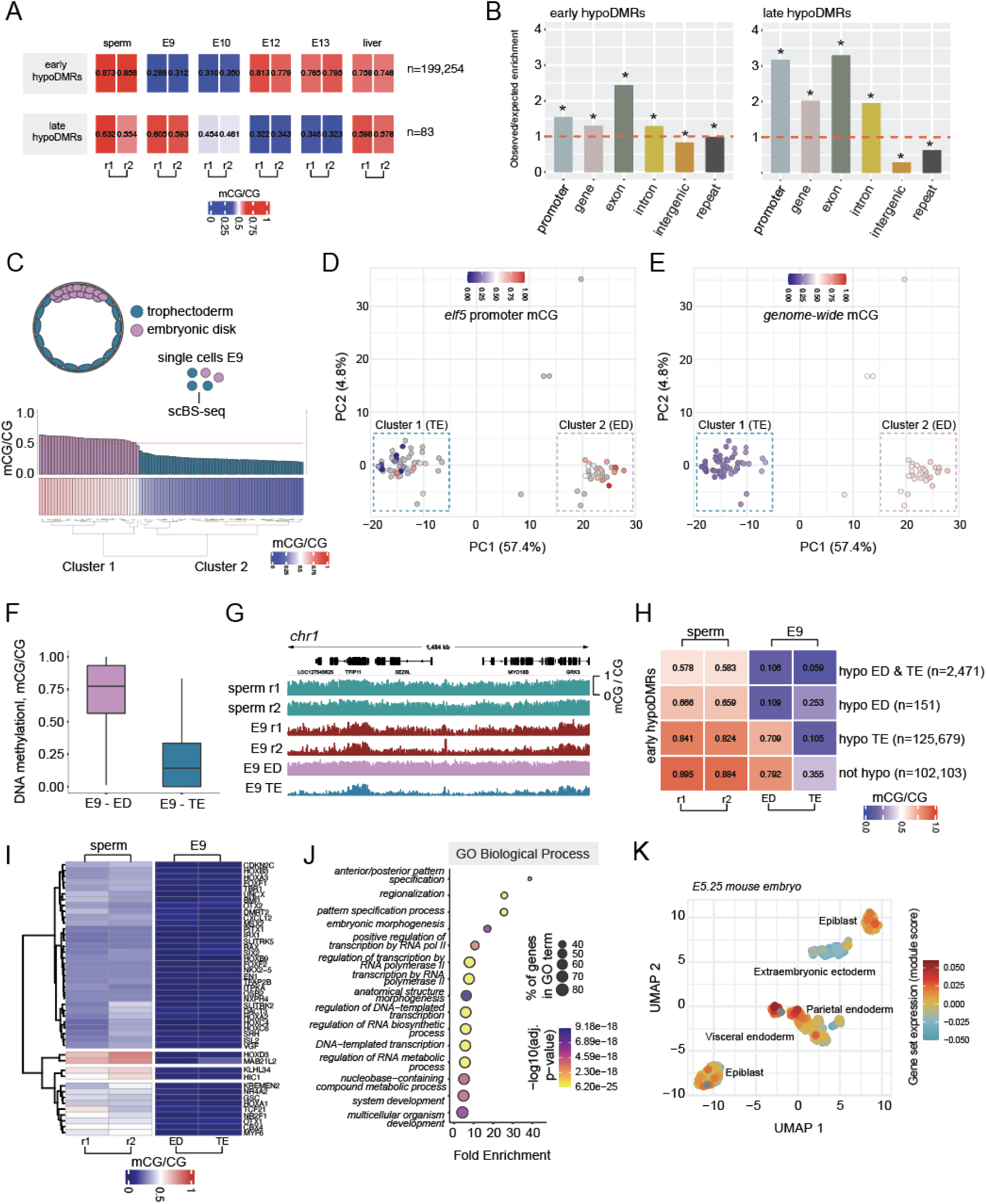
Single cell DNA methylation analysis of embryonic and extraembryonic lineages in the E9 dunnart embryo. **(A)** Heatmaps depicting two classes of hypomethylated DMRs in the dunnart embryo: early (hypomethylated in E9 embryos) and late (hypomethylated in E12 embryos). **(B)** Observed over expected enrichment plots showing an enrichment of early- and late hypoDMRs at different genomic regions. Asterisk indicates significant enrichment (p-value < 0.05). **(C)** Barplot and heatmap showing average genome methylation levels in single cells from E9 dunnart embryo. **(D)** Principle component analysis (PCA) of single cell promoter DNA methylation in the E9 dunnart embryo, depicting per-cell mean methylation of the trophectoderm marker gene promoter *Elf5*. Cells coloured in grey lack sufficient read coverage for the promoter of interest. **(E)** PCA of single cell promoter DNA methylation in the E9 dunnart embryo, depicting per-cell genome-wide mean methylation values. **(F)** Boxplots of genomic 5mC levels (1 kb non-overlapping genomic tiles) of pseudobulked E9 embryo trophectoderm (TE) and embryonic disk (ED). **(G)** IGV browser track depicting bulk (EM-seq) DNA methylation profiles of fat-tailed dunnart sperm and E9 embryos, as well as pseudobulked E9 embryo TE and ED. **(H)** Heatmap stratifying whole E9 embryo early hypoDMRs into four classes depending on the methylation levels in ED and TE. **(I)** Heatmap depicting gene body DNA methylation of genes that undergo DNA methylation reprogramming in ED. **(J)** Gene ontology (GO) analysis of the genes that undergo DNA methylation reprogramming in ED. Biological Process ontology results are shown. **(K)** Uniform manifold approximation and projection (UMAP) projection of 331 individual cells from the E5.25 mouse embryo. Module score of the gene set with DNA methylation reprogramming in ED is plotted for each cell. Module scores depict the difference between the average gene expression of the gene set and random control genes.

We identified two distinct clusters of cells based on the average genome methylation levels (Fig. 2C). In order to establish the cell type of each cluster, we performed PCA analysis of promoter DNA methylation (Supplementary Fig. 2A). We assigned each cluster to the TE and ED based on the promoter methylation of canonical lineage markers *Elf5* (hypomethylated in TE), and *Nanog* (hypomethylated in ED) (Fig. 2D, Supplementary Fig. 2B)^63,64^. Genome-wide DNA methylation analysis of pseudo-bulked TE and ED samples revealed significant DNA methylation erasure in TE (23.2% global 5mC) but not in the ED (60.7% global 5mC) highlighting that the bulk E9 methylation profiles are largely reflective of the methylation state of the TE (Fig. 2D-G, Supplementary Table 2). This is in striking contrast with the South American opossum which displays 46.4% mean global 5mC in the E7.5 TE^48^, revealing a surprising evolutionary divergence in 5mC reprogramming strategies specifically in extraembryonic lineages (Supplementary Table 2). In contrast, DNA methylation maintenance in the embryonic lineage remained conserved in the divergent Australian and American marsupial families, with 60.2% and 60.7% mean global 5mC in the opossum and dunnart respectively (Supplementary Table 2).

To explore whether any genomic regions undergo DNA methylation reprogramming in the embryonic lineage, we classified sperm-E9 DMRs based on the residual methylation in the embryonic lineage and TE (see Methods) (Fig. 2H). The analysis identified 2471 hypoDMRs in the embryonic lineage, mostly located in fully demethylated gene bodies (Fig. 2I, Supplementary Fig. 2C). These were enriched in transcription factor genes that are crucial for developmental processes such as anterior-posterior patterning and anatomical structure morphogenesis (Fig. 2J). These key developmental genes are positioned within DNA methylation valleys, that are large hypomethylated blocks conserved in vertebrate genomes^65,66^.

To investigate the functional significance of these DNA methylation changes, we performed gene expression analysis using publicly available E5.25 mouse single cell RNA-seq data (Supplementary Fig. 2D)^67^. Epiblast and extraembryonic ectoderm cell clusters were validated by the *Nanog* and *Elf5* gene expression patterns, respectively (Supplementary Fig. 2E). The analysis revealed that genes whose bodies display DNA methylation reprogramming are actively expressed in the mouse epiblast, a tissue that gives rise to the embryo proper; but remain silent in the extraembryonic ectoderm, a tissue derived from the TE that contributes to ectoplacental structures (Fig. 2K, Supplementary Fig. 2F). These findings indicate that developmental genes still undergo conserved 5mC erasure, revealing a locus-specific regulatory role for DNA methylation that is independent of global demethylation.

### Genome-wide DNA methylation erasure in trophectoderm occurs alongside retention at specific gene bodies

In contrast to the embryonic lineage, the TE of the E9 dunnart embryo is extensively demethylated (Fig. 2H) displaying low DNA methylation across the entire genome including gene bodies, intergenic regions and repetitive elements (Fig. 3A, Supplementary Fig 3A, B). To compare the extent of TE methylation erasure across species, we re-analysed mouse and opossum TE methylation data^48,68^. The level of residual DNA methylation in dunnart TE (23.2% mC) was similar to that of mouse TE (18%), but markedly different from the American marsupial grey short-tailed opossum (46%) (Supplementary Table 2). Given such striking differences, we expanded our analysis to look at a range of mammalian species using publicly available TE and placental methylomes^48,68–70^. The analysis revealed a striking diversity of global DNA methylation levels across mammals with mouse and dunnart TE exhibiting the lowest global DNA methylation (∼25%), while horse and squirrel monkey placentas displayed significantly higher DNA methylation (∼60%) (Fig. 3B). Given the great variety in trophoblast invasiveness across mammals^71^, we initially hypothesised that invasiveness might correlate with the extent of DNA methylation erasure in the trophectoderm. However, we found no correlation between DNA methylation levels and the degree of placental invasiveness as classified by interhaemal barrier type (hemochorial, endotheliochorial, epitheliochorial) (Fig. 3B). This lack of correlation is consistent with placental invasiveness being controlled primarily by the maternal endometrium rather than trophoblast characteristics, since ectopic pregnancies demonstrate that trophoblasts possess intrinsically invasive capacity when unconstrained by endometrial regulation^72–74^ (Fig. 3B).

**Figure 3:**
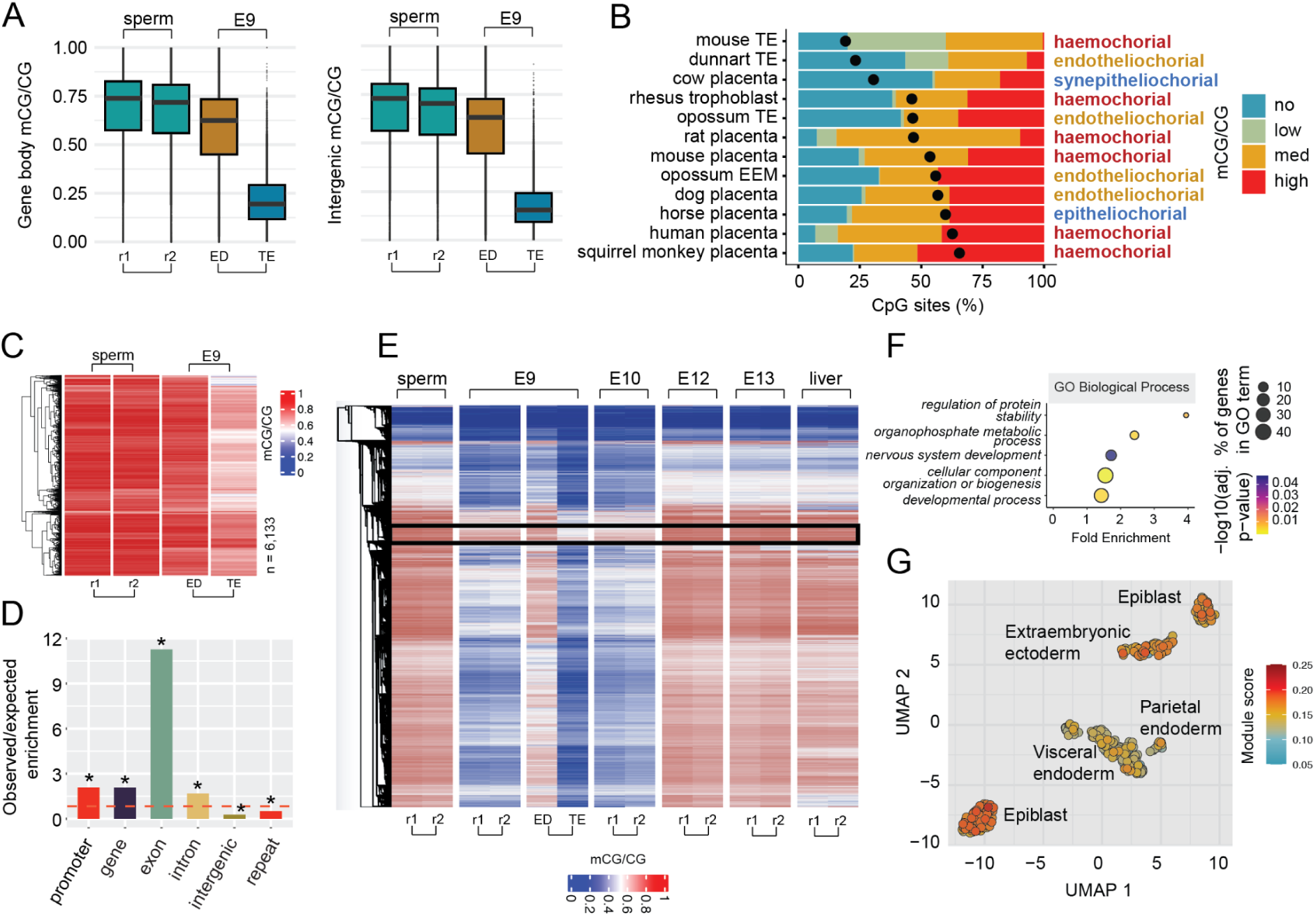
Trophectoderm DNA methylation landscape in fat-tailed dunnart E9 embryos. **(A)** Boxplots showing average gene body and intergenic regions DNA methylation levels in sperm and E9 embryo trophectoderm (TE) and embryonic disk (ED). **(B)** Stacked bar plots indicating the percentages of methylated CpG sites in trophectoderm (TE), trophoblast, extraembryonic membrane (EEM) and placental tissue across eutherian and metatherian species. High, 80-100%; medium, 20-80%; low, >0-20%; no, 0% methylation. Black dots indicate genome-average DNA methylation levels. Interhaemal barrier type (hemochorial, endotheliochorial, epitheliochorial) is colour-coded by the degree of invasiveness from high (red), to medium (yellow), to low (blue)^48,68–70^. **(C)** A heatmap depicting DNA methylation profiles of ‘invariant regions’ that retain high levels of DNA methylation in TE. **(D)** Observed over expected enrichment plot showing an enrichment of ‘invariant regions’ with high DNA methylation in TE at different genomic regions. **(E)** A heatmap of gene body DNA methylation profiles across dunnart development. Genes retaining high methylation at their gene bodies are highlighted. **(F)** Gene ontology (GO) analysis of the genes that retain high methylation levels in TE. Biological Process ontology results are shown. **(G)** Uniform manifold approximation and projection (UMAP) projection of 331 individual cells from the E5.25 mouse embryo. Module score of the gene set with high DNA methylation in TE is plotted for each cell. Module scores depict the difference between the average gene expression of the gene set and random control genes.

Despite genome-wide DNA methylation erasure in the TE, we observed that some genomic regions retained high levels of methylation. To explore them in more detail, we called ‘invariant’ CpG dinucleotides that retain high levels of methylation (0.7-1.0 methylated CG/total CG) throughout development and grouped adjacent sites into ‘invariant regions’ (Fig. 3C, n=6133). Observed over expected enrichment analysis revealed a significant over-representation of constitutively highly methylated regions among exons and gene bodies (Fig. 3D). To explore the retention of 5mC at gene bodies in more detail, we calculated average methylation per gene with hierarchical clustering, revealing a group of genes retaining high 5mC throughout development (Fig. 3E). Gene ontology analysis revealed an enrichment of genes involved in fundamental cellular processes and embryonic development (Fig. 3F, Supplementary Fig. 3C). Notably, genes with high gene body 5mC showed expression across both embryonic and extra-embryonic lineages in mouse E5.25 embryos (Fig. 3G, Supplementary Fig. 3D) in line with previous findings^69^. In contrast, genes that underwent 5mC erasure exhibited substantially reduced embryonic expression (Supplementary Fig. 3E), suggesting the deposition of DNA methylation at actively transcribed gene bodies^75–79^.

Finally, we sought to explore DNA demethylation dynamics at repetitive elements given that certain classes of transposable elements are known to retain higher residual DNA methylation during pre-implantation development in eutherian mammals^18,19^. The analysis showed that DNA methylation was maintained across various repeat classes in the embryonic lineage, while it was erased in the extraembryonic lineage (Supplementary Fig. 3F).

### Paternal X chromosome exhibits global hypomethylation except at escapee loci

Finally, we leveraged our new chromosome-level genome assembly to investigate epigenetic regulation of the inactive X chromosome in the fat-tailed dunnart. Given that our embryonic EM-seq data was generated from pooled embryos of undetermined sex, we profiled DNA methylation using long-read sequencing of adult male and female liver tissue. The analysis showed that methylation levels on the single X chromosome in males (60.7%) were nearly double those observed on the X chromosomes in females (37.5%) (Fig. 4A, B), suggesting a potential heterogeneity between maternal and paternal X chromosomes in the female sample. To address this, we called heterozygous germline SNPs to segregate two haplotypes using pipeface v1.0 (see Methods). Notably, only part of the X chromosome had enough heterozygosity to phase the two haplotypes, which is expected given the inbred nature of the dunnart colony (Fig. 4A). Next, to determine which of the two haplotypes corresponds to the maternal allele and which is paternal, we performed long read RNA-seq of cerebellum from the same female individual and segregated transcriptomic reads between haplotypes using GATK ASEReadCounter. The analysis revealed that the majority of transcripts correspond to haplotype 1, suggesting that it was an active, and therefore maternal, X chromosome (Fig. 4C). In contrast, haplotype 2 had lower expression, suggesting that it was an inactive, paternal X chromosome (Fig. 4C). Haplotype-resolved DNA methylation analysis of the two X chromosomes in females revealed that the active, maternal X (Xm) chromosome was hypermethylated (mean methylation=62.0%), while the inactive, paternal X (Xp) chromosome was hypomethylated (mean methylation=13.7%) (Fig. 4B, C), in line with recent findings in the opossum^48^ and koala^80^. Furthermore, in contrast to eutherians, gene promoters on the dunnart’s inactive X chromosome remained hypomethylated, consistent with observations in other marsupials (Supplementary Fig. 4A)^81–83^. This further supports that DNA hypermethylation as a repressive mechanism in X chromosome inactivation (XCI) is an evolutionary innovation unique to eutherians.

**Figure 4:**
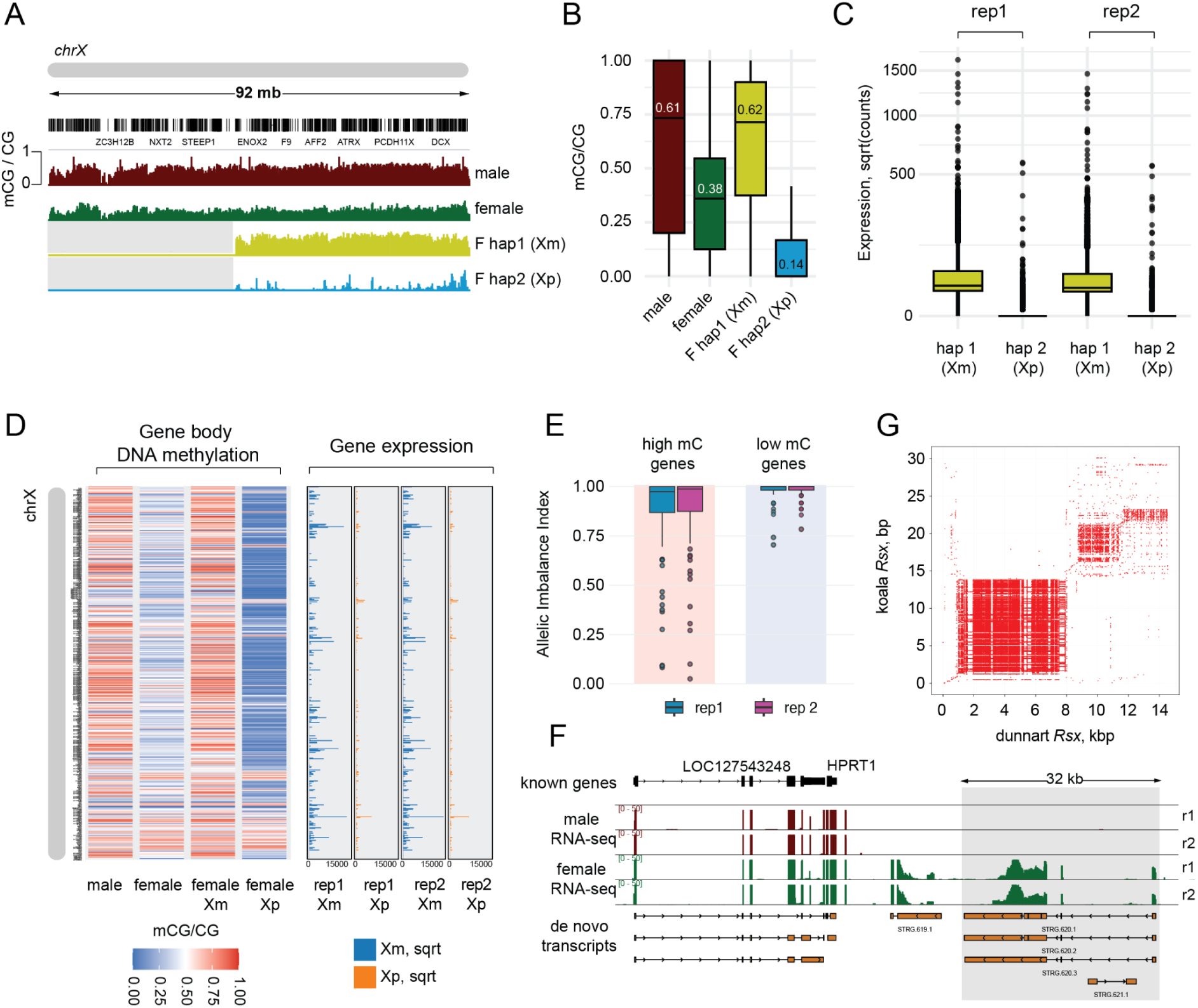
Epigenetic and transcriptional profiling of maternal and paternal X chromosomes. **(A)** IGV browser track depicting DNA methylation profiles of the X chromosome in male and female dunnart liver tissue. DNA methylation profiles of two female X chromosomes were further separated using haplotype-specific DNA methylation analysis. **(B)** Boxplots of genomic 5mC levels (1 kb non-overlapping genomic tiles) of male and female X chromosomes. Female X chromosomes are further separated into maternal (Xm) and paternal (Xp). **(C)** Boxplots showing expression of X-linked genes on Xm and Xp in female cerebellum. **(D)** Heatmaps showing haplotype-resolved DNA methylation and gene expression profiling along the length of the X chromosome. The distal part of the X chromosome (52Mb) that displays sufficient heterozygosity is shown (not greyed-out area on IGV track in panel A). **(E)** Allelic imbalance, AI = 2 * abs(0.5 – (Xm/Xm+Xp)), of the genes with high vs low gene body methylation on the paternal X chromosome. **(F)** IGV browser visualisation of the lncRNA *Rsx* locus (grey shading) showing RNA-seq data from male and female dunnart cerebellum, annotated genes, and *de novo* assembled transcriptome. **(G)** Dot plots comparing the conservation of repeat domains of the fat-tailed dunnart and koala lncRNA *Rsx*.

We noticed, however, that there were hypermethylated foci on the hypomethylated paternal X chromosome concentrated towards the distal end of the chromosome (Fig. 4D) spanning gene bodies (Supplementary Fig. 4B). To assess functional significance, we grouped adjacent highly methylated CpGs (0.7–1.0 methylated CG/total CG) on the paternal X chromosome into highly methylated regions and identified overlapping genes. Allele-specific expression analysis showed that these highly methylated genes exhibit reduced Xm/Xp allelic imbalance compared with hypomethylated genes on the paternal X (Fig. 4E). Reduced Xm/Xp imbalance reflects escape from XCI, indicating expression of highly methylated genes from the inactive Xp. Moreover, orthologues of the highly methylated dunnart genes were enriched, although not significantly, among known human escapee genes (Supplementary Fig. 4C). These findings suggest that rare gene-body DNA hypermethylation on Xi is a consequence of transcriptional activity, likely mediated by DNMT3 recruitment to H3K36me3-marked regions, similar to the mechanism observed in eutherians^75–78^.

Finally, we sought to identify the lncRNA *Rsx*, a known master regulator of X chromosome inactivation in marsupials, within the fat-tailed dunnart genome^50^. Given the absence of *Rsx* annotation in the publicly available dunnart GTF file, we performed reference-guided transcriptome assembly of female cerebellum long-read RNA-seq using *stringtie*^84^. We searched for female-specific *de novo* transcripts in a conserved syntenic region harbouring *Phf6* and *Hprt1* genes on the X chromosome^50^. The analysis identified a novel 32kb long gene containing 6 exons, 21 kb downstream from *Hprt1*, specifically expressed in female tissue (Fig. 4F). Comparative sequence analysis between dunnart and koala *Rsx* orthologs revealed extensive homology between their repeat domains (Fig. 4G), similar to that between koala and opossum *Rsx*^85^. BLASTN analysis identified multiple alignments ranging from large continuous segments at the 5’ region (up to 7.4 kb at 65% identity) to numerous shorter high-identity fragments at the 3’ region (70-95% identity), reflecting the tandem repeat structure of *Rsx* (Supplementary Table 4)^86^. The repetitive core generated hundreds of overlapping alignments, with individual dunnart segments matching multiple positions in the koala sequence, consistent with independent repeat unit evolution between species. Notably, the spliced dunnart *Rsx* transcript (14.5 kb) was shorter than the koala *Rsx* (23 kb), suggesting differential repeat expansion between these marsupial lineages. Overall, despite moderate overall identity (65%), the highly significant E-values and bit scores confirm high *Rsx* sequence orthology, pointing to functional conservation of the master regulator of X-chromosome inactivation across marsupial evolution.

Taken together, these findings highlight the conservation of DNA methylation landscapes on active and inactive X chromosomes in female marsupials, as well as lncRNA *Rsx* tandem repeat domains in X chromosome inactivation.

## DISCUSSION

5mC represents one of the most abundant epigenetic modifications in eukaryotic genomes, with clearly established roles in developmental gene regulation across vertebrates^1,3^. However, our current understanding of 5mC reprogramming in non-eutherian mammals is strikingly limited, with only one marsupial species, the South American opossum (*Monodelphis domestica*), being profiled to date^47,48^. The aim of this study was to explore how conserved or divergent 5mC reprogramming strategies are among therian mammals, and what this reveals about the evolutionary significance of this epigenetic process. Our near-complete T2T genome assembly and base-resolution 5mC maps of embryonic development in the fat-tailed dunnart revealed that the dunnart retains a hypermethylated embryonic disk in the bilaminar blastocyst, reflecting developmental epigenome dynamics that diverge from eutherians, yet are conserved across both Australian and American marsupials. However, the dunnart undergoes extensive 5mC loss in the trophectoderm, pointing towards deep evolutionary conservation in the epigenetic regulation and development of placental tissues across mammalian lineages.

The absence of global 5mC erasure in the marsupial embryonic lineage prompts a fundamental question: what is the functional significance of this process in eutherian mammals? It is speculated that 5mC erasure in eutherian lineages is linked to the early onset of zygotic genome activation (ZGA) in mammals. For example, there are ten and twelve cell divisions before ZGA in zebrafish and frog respectively, while in mice both minor and major waves of ZGA occur by the late two-cell stage^87^. The extended period prior to ZGA initiation in zebrafish may allow for fine-tuning of the embryonic epigenome through loci-specific 5mC changes, rather than by expeditious global DNA demethylation^20,21^. In support of this hypothesis, 5mC erasure in sheep and rabbit is less extensive than in mouse and human, which is correlated with a later onset of ZGA in these species^58^. However, in the dunnart bilaminar blastocyst (E9), and throughout cleavage stages in the opossum (E1.5 to E4.5), the embryonic genome maintains high methylation levels (>60%), contrasting sharply with the progressive demethylation seen in most eutherian species (Fig. 2C-G)^48^. This hypermethylated state persists even through embryonic genome activation at E3.5 in the opossum^88^, demonstrating that extensive demethylation is not required for this critical developmental milestone, at least in the marsupial species profiled. However, the distinction of early versus late ZGA onset distinction must be interpreted with reservation, as mammals undergo cleavages more slowly than non-mammalian vertebrates^89^, suggestive of a more complex relationship between post-fertilisation 5mC dynamics and ZGA among vertebrate species.

The fundamental differences in embryonic methylation dynamics across mammalian species may have important implications for the cellular reprogramming strategies. It was recently shown that transient-naïve-treatment (TNT) reprogramming of human somatic cells to iPSCs more faithfully recapitulates the epigenetic state of pre-implantation embryos^90^ compared to conventional OKSM or SCNT reprogramming^91^, by facilitating H3K9me3 heterochromatin remodelling and transient genome-wide DNA demethylation. Given that DNA methylation erasure varies from extensive (mice, humans) to moderate (sheep, rabbits, marsupials), testing TNT reprogramming across these species could reveal whether its efficacy in generating iPSCs correlates with natural embryonic DNA demethylation dynamics.

Despite divergent embryonic reprogramming strategies in eutherians and marsupials, we identify deep evolutionary conservation of trophectoderm 5mC erasure. It is plausible that, in eutherians, 5mC erasure in both the embryo proper and trophectoderm facilitates the early segregation of these lineages, which is necessary for markedly earlier implantation compared to marsupials^88,92^. In mice, which have a 21-day gestation period, the ICM becomes distinguishable from the trophectoderm by E3.5, just before implantation. In contrast, the opossum and dunnart do not exhibit clear segregation of embryonic and TE lineages until E6.5 and E9, respectively, despite having pregnancies lasting only 13 days^48,49,88^. Therefore, this slower developmental progression may provide the means for cell-type specific global DNA demethylation. Intriguingly, we found that 5mC levels in the dunnart trophectoderm are significantly lower compared to those in the opossum (Fig. 3B), suggesting evolutionary divergence in epigenetic regulation of the placental precursor among marsupials, mirroring patterns observed in eutherians^69^. Nevertheless, our findings confirm that 5mC erasure in the trophectoderm represents an ancestral state in therian mammals, and may have been important for the evolution of a trophoblast with the capacity to invade the uterine mucosa, evade the maternal immune system, and regulate the maternal/fetal interface.

It is unclear exactly why the placenta remains partially methylated during gestation. The functional significance of DNA demethylation in trophectoderm lineage restriction is evident from studies showing that DNA methyltransferase-deficient embryonic stem cells (ESCs) and embryos demonstrate enhanced trophoblast differentiation potential. Specifically, *Dnmt1*−/− ESCs showed a 5-fold increased contribution to trophectoderm compared to wild-type controls^63,93^. Similarly, in *Dnmt* triple knockout (TKO) and wild-type (WT) chimeric embryos, TKO cells were rarely found within the embryo proper but showed substantial contribution to extraembryonic lineages^94^. Furthermore, inhibiting DNA methyltransferases to prevent aberrant re-methylation in somatic cell nuclear transfer (SCNT) embryos, enhanced their developmental potential and reduced placental abnormalities in cloned embryos^4^. The unique 5mC landscape of the placenta is thought to facilitate the expression of genes related to angiogenesis, tissue invasion and immunosuppression, with analogous molecular and morphological features to many cancers^68,95,96^. This would indicate that diversity in global 5mC levels are linked to such phenotypes; indeed, dunnart and opossum placentas differ in the degree to which fetal tissues invade the maternal endometrium, which may partially explain these epigenetic discrepancies^97^.

While the majority of regions undergoing DNA methylation erasure in E9 dunnart embryos were demethylated in trophectoderm and methylated in the embryonic lineage, a smaller subset of regions (n=2471) also exhibited DNA methylation erasure in the embryonic lineage (Fig. 2H). These regions were associated with gene bodies of developmental genes including *Hoxa1*, *Pitx1*, and *Pax6* (Fig. 2I, J). This observation aligns with previous findings demonstrating that the majority of key developmental genes across different vertebrate species are located within large hypomethylated blocks, termed DNA methylation valleys (DMVs) and canyons^65,98^. This hypomethylated state has been suggested to confer protection against spontaneous deamination of 5mC and likely facilitates plasticity in gene expression due to the high density of transcription factor binding sites^66^.

Although this study is focused on embryonic 5mC reprogramming, it is worth noting that mammals have also evolved the ability to globally reset their methylome in primordial germ cells (PGCs), enabling the erasure and re-establishment of 5mC at imprinted genes. This would imply that there is no requirement for global 5mC erasure in species where these processes do not occur; indeed, zebrafish retain paternal 5mC patterns in PGCs^99,100^. In the brushtail possum (*Trichosurus vulpecula*) and tammar wallaby (*Macropus eugenii*), substantial primordial germ cell (PGC) DNA demethylation has been reported^46,101^, pointing to independent evolution of embryonic and germline DNA demethylation in mammals. However, it is important to note that to date there are no comparative datasets of both embryonic and germline 5mC reprogramming in a single marsupial species.

Finally, our T2T genome assembly enabled identification of i) the previously unannotated long non-coding RNA *Rsx* on the dunnart X chromosome, and ii) global hypomethylation of the paternally derived inactive X in adult female somatic tissue (Fig. 4). In the opossum and tammar wallaby, chromosome-wide 5mC levels on the X chromosome in sperm resembles that of autosomes^48,102^, indicating that the paternal inactive X is specifically targeted for global DNA hypomethylation during early embryogenesis. It is unlikely that this occurs via exclusion of the maintenance methyltransferase DNMT1 from the nucleus, as hypomethylation is specific to only one allele of one chromosome. Although it is possible that it is mediated by active DNA demethylation via Ten-Eleven Translocation (TET) enzymes, RNA-seq of opossum embryos revealed low *Tet* expression at the pronuclear stage^48^. In addition, 5mC at gene bodies is known to prevent spurious transcription initiation at cryptic promoters located within gene bodies^79^, indicating that hypermethylation is likely retained at XCI escapee genes through embryonic development. Future work may elucidate the mechanism by which the inactive X deters 5mC machinery during marsupial embryogenesis.

Taken together, our study offers the first detailed exploration of developmental 5mC dynamics in an Australian marsupial, greatly expanding our knowledge on the diversification of embryonic gene regulation across placental mammals, and how these processes contribute to the evolution of unique developmental strategies.

## SUPPLEMENTARY FIGURES

**Supplementary Figure 1:**
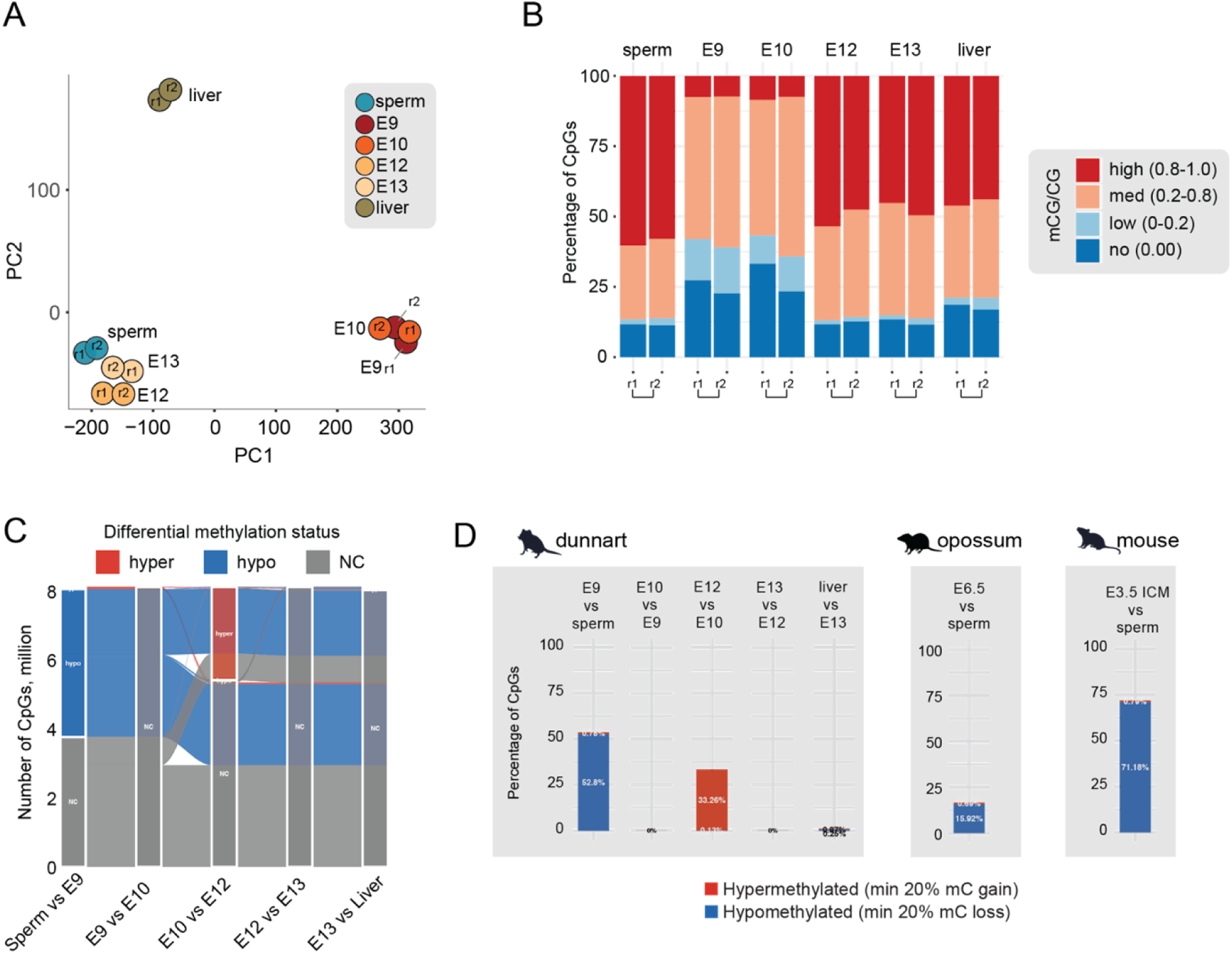
DNA methylation dynamics in the fat-tailed dunnart embryo. **(A)** Principal component analysis (PCA) of dunnart methylomes binned into 1kb regions. **(B)** Stacked bar plots indicating the percentages of methylated CpG sites in dunnart sperm, embryos and adult tissue. High, 80-100%; medium, 20-80%; low, >0-20%; no, 0% methylation. **(C)** Alluvial plot showing the number and dynamics of hypo- and hypermethylated differentially methylated cytosines (DMCs) between consecutive developmental stages. **(D)** Percentages of differentially methylated CpGs out of all covered CpGs in the genome between consecutive stages of embryonic development in the dunnart, opossum and mouse. CpGs with at least 20% methylation loss compared to the previous stage were considered as hypomethylated. CpGs with at least 20% methylation gain compared to the previous stage were considered as hypermethylated.

**Supplementary Figure 2:**
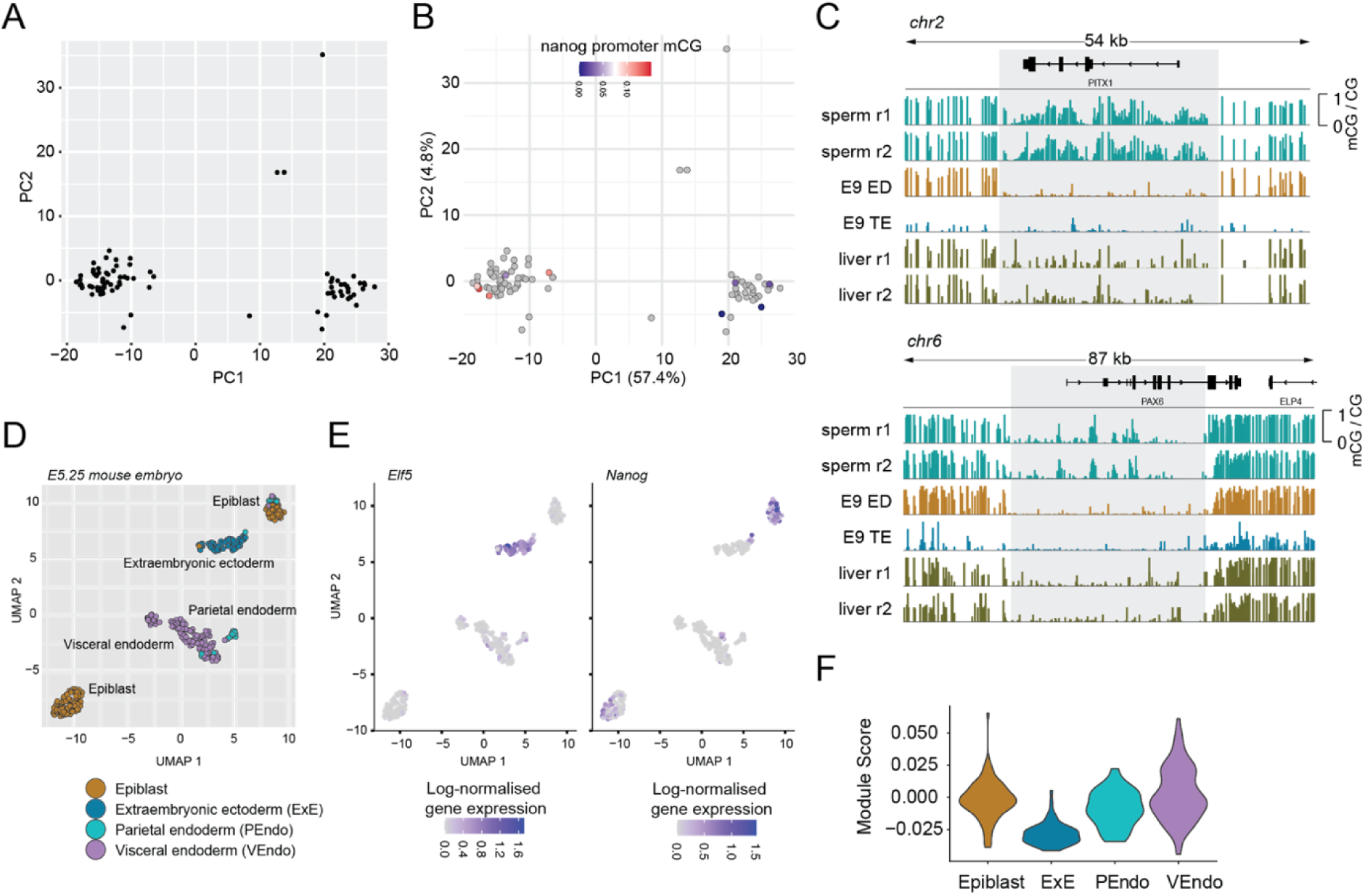
Single cell DNA methylation analysis of embryonic and extraembryonic lineages in the E9 dunnart embryo. **(A)** Principal component analysis (PCA) of single cell promoter DNA methylation in the E9 dunnart embryo. **(B)** PCA of single cell promoter DNA methylation in the E9 dunnart embryo, depicting per-cell mean methylation of the embryonic marker gene promoter *Nanog*. Cells coloured in grey lack sufficient read coverage for the promoter of interest. **(C)** IGV browser track depicting sperm and liver EM-seq as well as pseudobulked E9 embryo’s TE and ED scBS-seq DNA methylation profiles. **(D)** Uniform manifold approximation and projection (UMAP) of 331 individual cells from the E5.25 mouse embryo. Cells are color-coded by tissue types. **(E)** UMAP showing expression levels of the extraembryonic ectoderm marker *Elf5* and the epiblast marker *Nanog*. **(F)** Violin plots showing module scores of the gene set that undergoes DNA methylation reprogramming in embryonic and extraembryonic lineages. Module scores depict the difference between the average gene expression of the gene set and random control genes. ExE - extraembryonic ectoderm, PEndo - parietal endoderm, VEndo - visceral endoderm.

**Supplementary Figure 3:**
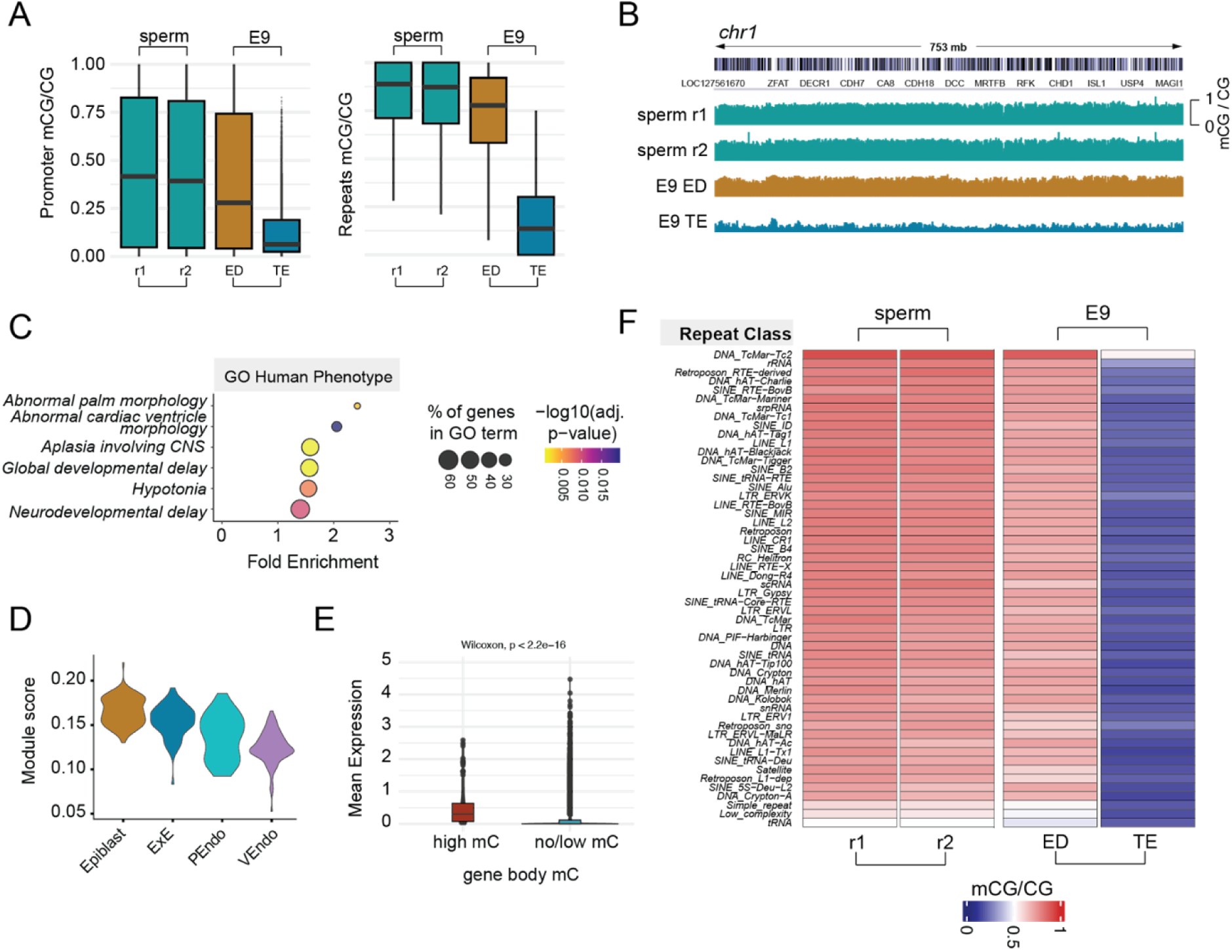
Genome-wide DNA methylation erasure in the dunnart’s trophectoderm. **(A)** Boxplots showing average promoter and repetitive elements DNA methylation levels in sperm and E9 embryo’s trophectoderm (TE) and embryonic disk (ED). **(B)** IGV browser track depicting sperm EM-seq and pseudobulked E9 embryo’s TE and ED scBS-seq DNA methylation profiles. The entire chr1 is shown. **(C)** Gene ontology (GO) analysis of the genes that retain high mC levels in TE. The human phenotype ontology results are shown. **(D)** Violin plots showing module scores of the gene set with high DNA methylation in TE. Module scores depict the difference between the average gene expression of the gene set and random control genes. ExE - extraembryonic ectoderm, PEndo - parietal endoderm, VEndo - visceral endoderm. **(E)** Boxplots showing mean expression levels of genes that retain high gene body DNA methylation vs genes that undergo DNA methylation erasure at their gene bodies. **(F)** Heatmap showing DNA methylation at transposable elements in sperm, E9 embryo’s trophectoderm (TE) and embryonic lineage (ED).

**Supplementary Figure 4:**
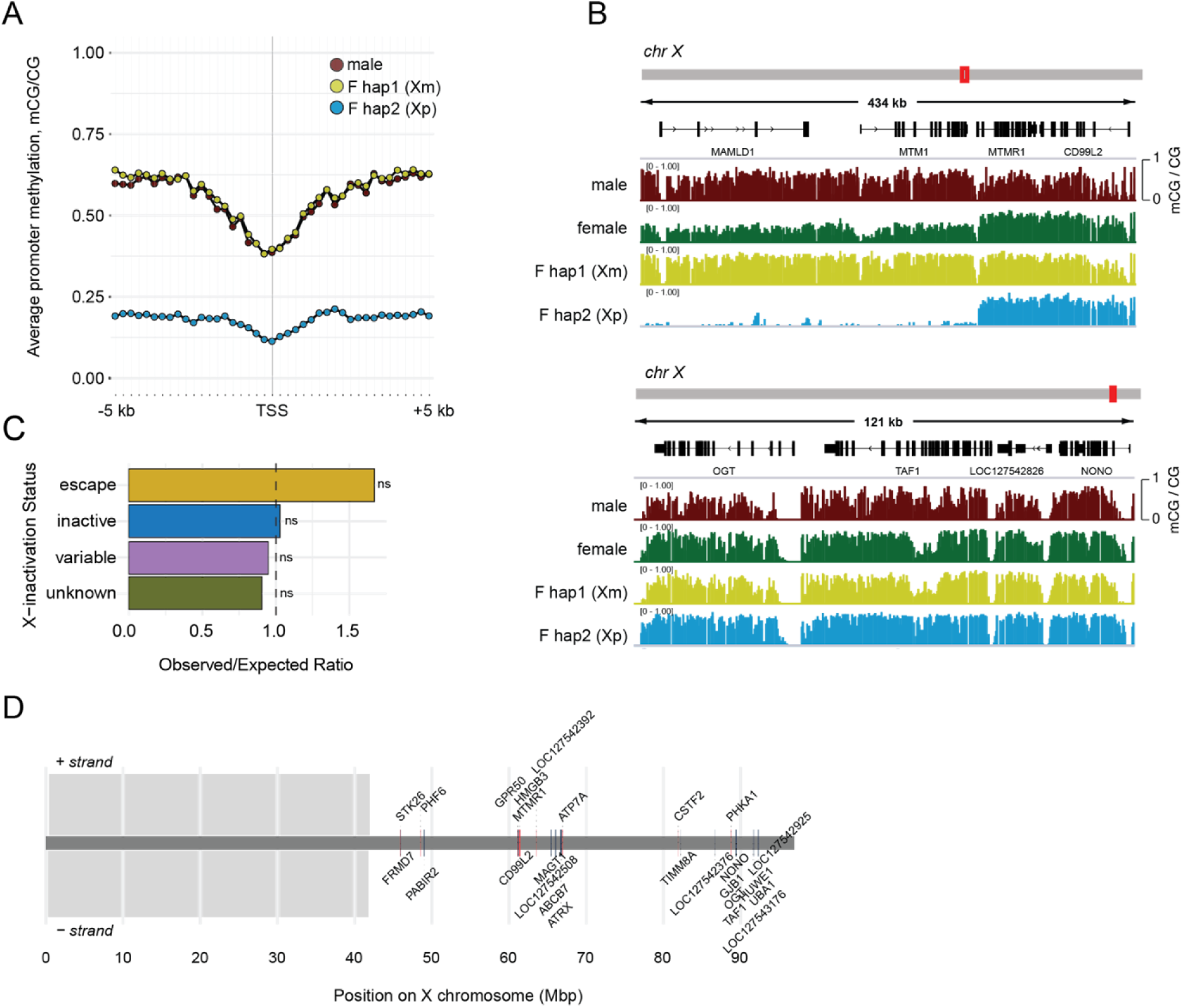
DNA methylation profiling of active and inactive X chromosomes and escape from X chromosome inactivation in the fat-tailed dunnart. **(A)** Promoter DNA methylation profiling of the active maternal (yellow) and inactive paternal X chromosomes in the female, as well as single male X chromosome (red) in the adult liver. **(B)** IGV browser track showing examples of genes that exhibit unusually high DNA methylation levels on the inactive paternal X chromosome in female dunnart liver. **(C)** Observed over expected enrichment of highly methylated dunnart genes on the inactive Xp among known human escapee, variable and inactive X-linked genes. **(D)** Location of putative fat-tailed dunnart escapee genes on the X chromosome.

## METHODS

### Collection of fat-tailed dunnart embryos, gametes and adult tissues

Tissues were collected from a long term breeding colony of fat-tailed dunnarts at Macquarie University. All animal work was conducted with approval from the Macquarie University Animal Ethics Committee (ARA: 2020/019). Staged embryos were collected by daily weighing of paired females through the breeding season to identify the weight changes associated with ovulation^103^. Once at the right day of gestation, females were humanely killed with an intra-peritoneal injection of Sodium Pentobarbitone. Adult tissues were collected opportunistically, either from the same pregnant females or during tissue collections for other projects. Following dissection, embryos and tissues were snap frozen in liquid nitrogen for downstream processing. For sperm collection, the cauda epididymis was dissected from adult male testes after culling. Small incisions were made and the tissue was gently pressed to release sperm. Sperm were rinsed and collected in PBS, then snap-frozen in liquid nitrogen for downstream processing.

### Oxford Nanopore genome sequencing library preparation

DNA was extracted from dunnart liver using the PacBio Nanobind PanDNA kit (PN: 103-260-000), following manufacturer’s instructions. For standard Nanopore library preparation, 4 ug of DNA was sheared at speed 29 using a Megaruptor. Samples were prepared using the Nanopore SQK-LSK114 kit and sequenced for 96 hours on a PromethION FLO-PRO114M flow cell, with washes performed every 24 hours.

### Genome assembly

Data was basecalled with Dorado v0.8.3 (super accuracy model v5.0.0) via slow5-dorado to enable SLOW5 data format input^104^. This was used as input to create an initial assembly with Hifiasm (v0.25.0-r726)^105,106^. To improve on this assembly, a second flow cell was run using ‘Cornetto’ selectively sequencing and adaptive genome assembly, enriching ONT data onto the challenging unsolved genome regions, as described recently^54^. To support haplotype phasing and chromosome scaffolding, Hi-C was performed using dunnart tissue from the same individual. Hippocampus samples from a male and female adult dunnart were prepared for HiC using the Arima-HiC+ kit (Arima High Coverage HiC protocol, Low Input: Fresh Frozen Animal Tissue) according to manufacturer instructions. Library construction was performed using the Arima High Coverage HiC library preparation module according to manufacturer instructions, and libraries were sequenced on the Illumina NovaSeq X. Using both the Oxford Nanopore and Hi-C fastq files as input; Hifiasm was run, including the dual-scaf option to improve contiguity. Scaffolding was then performed using HapHiC (v1.0.7) using the documentation’s recommended alignment and filtering settings^107^. The output was then reviewed and manually curated in Juicebox, to obtain the final chromosome sequences for each dunnart.

### gDNA extraction for EM-seq

gDNA was extracted in biological replicate from dunnart embryonic samples using the QIAamp DNA Investigator Kit according to manufacturer instructions, and dunnart adult liver tissue using the QIAGEN DNeasy Blood & Tissue Kit according to manufacturer instructions. gDNA was isolated from sperm using the QIAGEN DNeasy Blood & Tissue Kit according to manufacturer instructions with modifications. Briefly, snapfrozen sperm in 100μL PBS was incubated with equal volume of Buffer X2 (20 mM Tris·Cl pH 8.0, 20 mM EDTA, 200 mM NaCl, 4% SDS, 80mM DTT, 12.5μl/ml QIAGEN Proteinase K), for one hour at 56°C. 200μL of Buffer AL and 200μL 100% ethanol were added to the sample, followed by centrifugation at 8000rpm for 1 minute in a DNeasy Mini spin column. 500μL Buffer AW1 was added to the sample which was then centrifuged at 8000rpm for 1 minute. 500μL Buffer AW2 was added to the sample which was then centrifuged at 14000rpm for 3 minutes. Sperm gDNA was then eluted in Buffer AE.

### EM-seq library preparation

Whole-genome methylation libraries were generated using NEBNext Enzymatic Methyl-seq (New England Biolabs) according to manufacturer instructions^56^. For each sample, unmethylated lambda DNA and methylated pUC19 DNA were spiked-in at 0.5% and 0.025% gDNA w/w, respectively. gDNA was then sonicated to an average length of 300 base pairs in the Covaris M220 Focused Ultrasonicator using the following settings: peak incident power, 50 W; duty factor, 20%; cycles per burst, 200; treatment time, 75 s. Sonicated DNA was concentrated in a vacuum centrifuge concentrator to a final volume of 50 µL, followed by DNA end-repair and dA-tailing, adaptor ligation, TET2 oxidation of 5mC and 5hmC, APOBEC deamination of unmodified cytosine to uracil, and PCR amplification. Library size and consistency was determined by the Agilent 4200 Tapestation system and libraries were quantified using the KAPA Library Quantification Kit according to manufacturer instructions. Libraries were sequenced on the Illumina NovaSeq X Plus platform (150bp, paired-end), covering >80% of all reference CpG sites with a mean coverage >6X per sample, per biological replicate.

### EM-seq methylome data analysis

Sequenced reads in fastq format were trimmed using Trimmomatic (trimmomatic PE -threads 24 ILLUMINACLIP:TruSeq3-PE-2.fa:2:30:10 SLIDINGWINDOW:5:20 LEADING:5 TRAILING:5 MINLEN:50)^108^. The chromosome-level assembly containing the lambda and pUC19 DNA sequences was indexed using WALT makedb command followed by mapping of trimmed reads to the indexed genome (walt -sam -m 10 -t 24 -N 10000000 -L 2000)^109^. Mapped reads in SAM format were converted to BAM and indexed using SAMtools^110^. PCR and optical duplicates were removed using Picard Tools (MarkDuplicates REMOVE_DUPLICATES=true) (http://broadinstitute.github.io/picard). Genotype and methylation bias correction was performed using MethylDackel software (https://github.com/dpryan79/MethylDackel). Reads containing at least three non-converted CpH sites (CpA, CpC, CpT) were identified and filtered from libraries using SAMtools and Picard FilterSamReads. The number of methylated and unmethylated calls at each genomic CpG position were determined using MethylDackel (MethylDackel extract –mergeContext --minOppositeDepth 5 --maxVariantFrac 0.5 --OT 20, 130, 20, 130 --OB 20, 130, 20, 130). Only cytosines in the CpG dinucleotide context were retained for further analyses. Enzymatic conversion efficiency was estimated from the lambda and pUC19 spike-in controls.

### Isolation and sorting of nuclei from E9 embryos

Single nuclei from 3 snap frozen E9 embryos were isolated using The S2 Genomics Singulator Platform, with the Low Volume Nuclei Isolation V2 protocol according to manufacturer instructions. DAPI-positive nuclei were sorted in 96-well plates containing 2.5μL of Buffer RLT (QIAGEN) and 1U/µL RNAse inhibitor using BD FACSAria™ III Cell Sorter. Plates were immediately placed on dry ice, sealed, and stored at −80°C until library preparation.

### Single cell bisulfite sequencing library preparation

Single cell bisulfite sequencing (scBS-seq) libraries were prepared as described with minor modifications^111^. First, RNA was removed from single cell lysates using oligo-dT conjugated magnetic beads^112^. Genomic DNA was then purified using a 0.8X volume of AMPure XP beads (Beckman Coulter) and subjected to bisulfite conversion using the EZ Methylation Direct Kit (Zymo). The oligo used in preamplification was modified to be compatible with NEBNext index primers used later in the protocol, and to prevent self-amplification: /5SpC3/CTACACGACGCTCTTCCGATCTNNNNNN (IDT, HPLC purified). After completing 5 rounds of preamplification and exonuclease treatment, the first-strand synthesis products were cleaned-up using a 0.65X volume of AMPure XP beads. Adaptor 2 tagging was then performed, and a 0.65X volume of buffer from AMPure XP beads was used to purify the second-strand synthesis products. Library amplification was then performed using NEBNext Multiplex Oligos for Illumina (Dual Index Primers) and 1x NEBNext Q5 Ultra II Master Mix (New England Biolabs), as follows: 98°C for 30 s; 14 repeats of 98°C for 10 s, 65°C for 30 s, 65°C for 45 s; 65°C for 5 min; 4°C hold. Amplified scBS-seq libraries were cleaned-up using a 0.65X volume of buffer from AMPure XP beads, assessed using the TapeStation High Sensitivity D5000 ScreenTape assay (Agilent), and quantified using Qubit dsDNA High Sensitivity assay (Thermo Fisher Scientific). Indexed libraries were then pooled in equimolar amounts for sequencing on the Illumina NovaSeq X Plus platform (150bp, paired-end), with 1,711,806-5,953,096 covered reference CpG sites for all cells passing QC.

### Single nuclei methylome data analysis

Sequenced reads in fastq format were trimmed using TrimGalore (trim_galore --cores 4 --paired --clip_r1 6 --clip_r2 6) (https://github.com/FelixKrueger/TrimGalore) and read alignment to the chromosome-level assembly was performed using Bismark (bismark --non_directional --unmapped --multicore 2), with reads not meeting paired-end constraints mapped in single-end format^113^. Mapped reads in BAM format were deduplicated using Bismark (deduplicate_bismark --bam --multiple), and per-CpG methylation levels were called using Bismark bismark_methylation_extractor (--gzip –bedGraph --paired-end/--single-end) and bismark2bedGraph (default settings) functions. PCA of promoter 5mC (including *Elf5*, *Nanog* and genome-wide methylation levels) was performed using MethSCAn^114^. Briefly, methylation calls were prepared using the methscan prepare function with default settings. Low-quality nuclei were removed for downstream analysis (methscan filter --min-sites 1000000 --min-meth 10 --max-meth 70), with 86/87 total profiled nuclei being retained (Supplementary Table 3). Smoothed mean methylation along the genome was calculated using methscan smooth command with default settings. A matrix of methylation levels at promoters (TSS +/− 1kb) was generated using the methscan matrix command with default settings, and PCA was performed using mean shrunken residuals of methylation levels at promoters, including iterative imputation of values at regions lacking sufficient coverage. Methylation levels at the *Elf5* and *Nanog* promoters (TSS +/− 1kb) were calculated for nuclei where at least half of all CpG sites located in the promoter sequence were covered by at least one read. Based on PCA clustering, nuclei were pseudobulked for downstream analysis by combining read counts from sample bedGraph files generated in the bismark2bedGraph command.

### DMR analysis

Analysis of differentially methylated CpGs (DMCs) and regions (DMRs) was performed using DMRcate^61,115^. Sequencing.annotate function was used to generate per-CpG t-statistics, indexing the FDR at 0.05. For the DMC analysis, a CpG site was considered significantly hypo- or hypermethylated between consecutive developmental stages, if the FDR < 0.05 and abs(delta mC) < 0.2 or abs(delta mC) < 0.5. The DMR calling was performed using DMRcate with minimum 5 CpGs and min.smoothed.fdr < 0.05. To classify sperm-E9 hypoDMRs as hypomethylated in ED, TE, or both lineages, we pseudobulked methylation levels for each lineage and calculated methylation differences (Δ) between sperm vs. ED and sperm vs. TE for each hypoDMR. We classified the DMR as hypomethylated in each lineage if the residual methylation in ED (or TE) was <=0.2 and Δ between sperm vs. ED (or sperm vs. TE) was <-0.3.

### Invariant CpG analysis

CpG sites with the methylation levels within the range [0.7, 1] mCG/CG and median coverage of 3 across sperm and embryonic samples were deemed as invariantly ‘highly methylated’. CpG sites located within 500 bp from each other were joined into invariant regions, which were filtered to harbor at least three invariant highly methylated CpGs.

### Repeat annotation

Repeats from the diploid male dunnart genome were analysed using RepeatMasker v4.2.0 with the settings “-no_is -species mammals”^116^. The RepeatMasker installation’s default engine was used, namely nhmmscan v3.4 (http://eddylab.org/software/hmmer/hmmer.org) (TRF v4.09.1^117^ as a dependency), and the HMM-Dfam v3.9 database^118^. To facilitate efficient use of computing resources, each chromosome was split into 15Mbp chunks (with 1.5kbp overlaps between chunks). The masked chunks were recombined using the bases furthest from the break points.

### Identification of mouse and human gene orthologues

To identify orthologues, amino acid sequences of protein-coding genes in fasta format of four eutherian species (*Homo sapiens* - GRCh38, *Mus musculus* - GRCm39, *Oryctolagus cuniculus* - OryCun2.0, *Sus scrofa* - Sscrofa11.1) and three metatherian species (*Monodelphis domestica* - ASM229v1, *Phascolarctos cinereus* - phaCin_unsw_v4.1, *Sarcophilus harrisii* - mSarHar1.11) were downloaded and filtered for the longest transcript variant per gene (OrthoFinder primary_transcript.py). The orthofinder command was run on dunnart and related mammalian species amino acid fasta files using default settings to identify mouse and human gene orthologues^119^.

### Mouse scRNA-seq data analysis

Mouse E5.25 embryo scRNA-seq^67^ Seurat object data was downloaded from https://tome.gs.washington.edu/. Read counts were normalized to 10,000 counts per cell and log-transformed. Highly variable features were identified with FindVariableFeatures, and UMI counts per cell were regressed out using ScaleData. Principal component analysis was performed on scaled data (RunPCA), followed by cell clustering using FindNeighbors (dims = 1:30) and FindClusters (resolution = 0.5, algorithm = 1). UMAP dimensionality reduction was applied using RunUMAP (dims = 1:30). Module scores were calculated using AddModuleScore for two gene sets: (1) genes whose gene bodies erase DNA methylation in the embryonic lineage and (2) genes whose gene bodies retain DNA methylation in trophectoderm.

### Gene annotation

*Sminthopsis crassicaudata* genes from the publicly available genome assembly were mapped to the new chromosome-level genome assembly generated in this study, using a lift-over tool liftoff^120^: liftoff -g ${OLD_GTF_DIR}dunnart_old.gtf $dunnart_male_T2T_21082025.fasta $Dunnart_asm_12-2021_sm.fa -o dunnart_male_T2T.gtf -u dunnart_male_T2T_unmapped.gtf -p 8 -copies -polish -exclude_partial -a 0.95 -s 0.95 -f transcripts.

### RNA extraction and cDNA sequencing

RNA was extracted from dunnart cerebellum using the QIAGEN RNeasy Mini Kit according to manufacturer instructions with DNAse treatment. PCR-cDNA library preparations were performed in technical replicates on each sample using the Nanopore SQK-PCB114.24 kit. The four barcoded amplified cDNA products were then pooled and sequenced on one PromethION flow cell (FLO-PRO114M) for 72 hours.

### Variant calling

The female X-chromosome scaffold was aligned to the male X reference with minimap2 (v2.24; preset asm5)^121^. Variants were called directly from the assembly to reference alignments using dipcall (v0.3) to produce a phased VCF. Long-read alignments from the same individual were then haplotagged with whatshap (v1.7) using this VCF^122^, and reads were split into hap1 and hap2. The haplotagged BAMs were processed with minimod to obtain haplotype-specific methylation calls.

### Oxford Nanopore RNA-seq data analysis

Male and female adult cerebellum long read RNA-seq was mapped using minimap2 using the following parameters: minimap2 -ax splice ${FASTA_DIR}$fasta ${FASTQ_DIR}$input > ${OUTPUT_DIR}$output_sam. Sam files were converted to bam files, sorted and indexed using SAMtools^110^. Duplicates were marked using Picard Tools MarkDuplicates (http://broadinstitute.github.io/picard). Read group information was added using Picard Tools AddOrReplaceReadGroups. To calculate allele-specific gene expression counts (refCount and altCount) for the X-linked genes, we applied gatk SplitNCigarReads followed by gatk ASEReadCounter -R ${FASTA_DIR}$fasta -I $input_bam -V ${VCF}femaletomaleX.dip_het_bi_PASS.vcf.gz -O $output --min-depth 10 --min-mapping-quality 20 --min-base-quality 20. Biallelic heterozygous variants were extracted from the vcf file using: bcftools view -m2 -M2 -g het $VCF -Oz -o femaletomaleX.dip_het_bi_PASS.vcf.gz. Allelic imbalance score was calculated using the following function: AI <-2*abs(0.5 - (refCount) / (refCount + altCount)).

### Reference-guided transcriptome assembly to identify lncRNA *Rsx*

First, long-read RNA-seq fastq files were aligned to the new reference genome generated in this study using minimap2 with the ‘-splice’ option as described above. Reference-guided transcriptome assembly of the female cerebellum long read RNA-seq was performed using stringtie: “stringtie ${BAM_DIR}$bam -G ${GTF_DIR}$gtf -o ${GTF_DIR}$denovo_gtf -l STRG“^84^. The syntenic region harbouring *Phf6* and *Hprt1* genes was visually scanned to locate stringtie-identified novel transcripts, expressed exclusively in female tissue. Dot plot analysis was performed in https://en.vectorbuilder.com/tool/sequence-dot-plot.html using default parameters (window size 10, mismatch limit 0).

## DATA AVAILABILITY

HiC, and ONT WGS and cDNA raw sequencing data generated for this study has been deposited at the European Nucleotide Archive (ENA) under accession number PRJEB98693. Enzymatic-methyl sequencing raw and processed data generated for this study has been deposited to ArrayExpress under accession number E-MTAB-16076. Single cell bisulfite sequencing raw and processed data generated for this study has been deposited to ArrayExpress under accession number E-MTAB-16075. Public datasets used in this study of the opossum, mouse and monkey embryonic and adult tissue BS-seq data are available under GSE206499, GSE56697, GSE60166, GSE146797 and GSE77124. Public datasets used in this study of trophectoderm and placental BS-seq datasets are available under GSE206499, GSE63330, GSE84235 and GSE211194. Public scRNA-seq data of the mouse E5.25 embryo are available under GSE109071.

## SUPPLEMENTARY TABLES

**Supplementary Table 1** Sequencing and enzymatic conversion quality metrics of EM-seq.

**Supplementary Table 2:** Average CpG methylation levels across the genome at various developmental stages in fat-tailed dunnart, opossum, mouse, and monkey embryos.

**Supplementary Table 3** Sequencing quality metrics of scBS-seq.

**Supplementary Table 4** Blastn alignment results comparing lncRNA *Rsx* sequences from fat-tailed dunnart and koala.

## CODE AVAILABILITY

Code used for processing the raw data, downstream analysis and the generation of figures is available on GitHub at https://github.com/kseskv/dunnart_mC_2025

## CONTRIBUTIONS

K.S., A.A. and O.W.G. conceived and designed the project. K.S. and A.A. wrote the manuscript. O.W.G. maintained the fat-tailed dunnart colony, collected embryos, and provided marsupial embryogenesis and placenta evolution expertise. A.A. performed EM-seq on dunnart embryos. A.A. aligned raw EM-seq and scBS-seq data. K.S. and A.A. performed computational data analysis. A.S. assisted with sequencing data quality control. T.P. performed DMR calling. V.P.M., M.W., K.C.K.I., K.P., R.L. and P.T. assisted with the embryo collection and single cell isolation. J.H., H.G., L.K. performed ONT dunnart genome and transcriptome sequencing, assembly and haplotype calling. A.L.M.R. and T.A. performed repeat annotation. A.D.S. performed mapping of publicly available DNA methylation datasets and provided help with mining the transferase database. E.D.W., N.L. and J.M.P. provided assistance with the genome annotation and fat-tailed dunnart lineage specification aspects. L.A.W., S.P. and L.J.R. performed dunnart sperm collection and DNA extraction. S.J.C. provided advice on the ONT methylation analysis of the X chromosome. H.L. and S.H. performed dunnart embryo scBS-seq library preparation. P.W. provided assistance with the *Rsx* gene annotation. I.W.D. provided advice on the ONT genome sequencing, chromosome-level genome assembly and allele-specific gene expression analysis. K.S. and O.W.G acquired funding for the project. All authors discussed the results and commented on the manuscript.

## AUTHOR INFORMATION

These authors contributed equally: Ksenia Skvortsova, Oliver W Griffith

## ACKNOWLEDGEMENTS

Single nuclei isolation and FACS (Fluorescence-activated cell sorting) services were provided by the Garvan Genomics Platform at the Garvan Institute of Medical Research. Specifically, we would like to thank Chris O’Keeffe, Eric Lam, Alejandro Rios Villamil, Samira Samarfard and Yasmin Husaini. Additionally, we are grateful to the Biomolecular Resources Facility at the Australian National University, and specifically to Maxim Nekrasov, for providing HiC-seq services. We are also grateful to the Garvan Molecular Genetics team, and specifically to Pavel Bitter and Chen Feng for low input genomic DNA extraction from the dunnart embryos. We are grateful to the Ramaciotti Centre for Genomics for providing EM-seq and scBS-seq sequencing services. We are grateful to Ozren Bogdanovic for critical reading of the manuscript and constructive feedback on the project. The research was supported by the Australian Research Council Discovery Project grant (DP250102459), National Health and Medical Research Council (2018114) (K.S.) and Discovery Project grant (DP200100344) and ARC Future Fellowship (FT240100181) (O.W.G).

